# Fungal States of Minds

**DOI:** 10.1101/2022.04.03.486900

**Authors:** Andrew Adamatzky, Jordi Vallverdu, Antoni Gandia, Alessandro Chiolerio, Oscar Castro, Gordana Dodig-Crnkovic

## Abstract

Fungal organisms can perceive the outer world in a way similar to what animals sense. Does that mean that they have full awareness of their environment and themselves? Is a fungus a conscious entity? In laboratory experiments we found that fungi produce patterns of electrical activity, similar to neurons. There are low and high frequency oscillations and convoys of spike trains. The neural-like electrical activity is yet another manifestation of the fungal intelligence. In this paper we discuss fungal cognitive capabilities and intelligence in evolutionary perspective, and question whether fungi are conscious and what does fungal consciousness mean, considering their exhibiting of complex behaviours, a wide spectrum of sensory abilities, learning, memory and decision making. We overview experimental evidences of consciousness found in fungi. Our conclusions allow us to give a positive answer to the important research questions of fungal cognition, intelligence and forms of consciousness.

## 1. The Basic Nature of Fungi

Cognition and intelligence in nature is a topic of debate from multiple points of view. While nowadays it is acceptable to speak about the cognition and intelligence within the biological kingdom Animalia, and even up to certain degree in Plantæ[80, 23], it is still controversial to discuss those capacities in other lifeforms such as Fungi, Protista and Monera, the latest corresponding to single-celled organisms without true nucleus (particularly bacteria) [85, 57]. However, the current move in research towards basal cognition and intelligence shows how already unicellular organisms possess basal levels of cognition and intelligent behaviour [43, 44, 45, 51, 52, 16, 11]. Further perplexity arises when considering consciousness in living organisms. Humans are conscious, and some allow consciousness in animals provided by a nervous systems. But other living creatures are typically considered not having consciousness at all. Our focus is on fungi, remarkable organisms with surprising cognitive capacities and behaviours which can be characterised as intelligent, and this article will argue that they possess a level of basal consciousness.

Fungi dominated the Earth 600 million years before the arrival of plants [84, 20]. Even today the largest known living organism in the world is a contiguous colony of *Armillaria ostoyae*, found in the Oregon Malheur National Forest, and colloquially known as the “Humongous fungus”. Its size is impressive: 910 hectares, possibly weighing as much as 35,000 tons and having an estimated age of 8,650 years [71]. Furthermore, current estimations sum up a total of 3.8 million existing species of fungi, out of which only 120.000 are currently identified [35], representing a promising biotechnological tool-set from which human kind has slightly scratched the surface.

Fungi have represented for humans ever since both an ally and a foe, in the first case serving to produce fermented food and beverages (just to cite the most important one, *Saccharomyces cerevisiae*, fundamental for bread, beer and wine) and in the second case, able to attack the same raw stocks and generate famine and devastation (*Puccinia graminis* responsible for stem, black or cereal rust) [29]. They have also shown particular features, including interaction with the nervous system of parasitised superior organisms, to induce them performing actions which are instrumental to further fungi propagation. This is the case of *Ophiocordyceps sinensis*, also known with its Tibetan name *yartsa gumbu*, an enthomopathogenic fungus parasitising ghost moths larvae, that is able to induce them standing vertically under the soil surface, to facilitate spores spreading in spring times [86]. Similarly, *Ophiocordyceps unilateralis* a complex of species also known as “zombie ant fungus” – surrounds muscle fibres inside the ant’s body, and fungal cells form a network used to collectively control the host behaviour, keeping the brain operative and guiding the ants to the highest points of the forest canopy, the perfect place to sporulate [55].

By studying Fungi kingdom we can better hope to understand the origin of life[60] and evolution of cognition, intelligence and consciousness as they gradually emerge from basal forms and up. But, beyond all the extremely important biochemical mechanisms that make them possible, a fundamental aspect in their organisation emerges: consciousness.

## 2. Neuron-like spiking of fungi

Spikes of electrical potential are an essential characteristic of neural activity [47, 14, 68]. Fungi exhibit trains of action-potential like spikes, detectable by intra- and extra-cellular recordings [75, 63, 2]. In experiments with recording of electrical potential of oyster fungi *Pleurotus djamor* (Fig. 1(a)) we discovered a wide range of spiking activity (Fig. 1(b)). Two types were predominant high-frequency (period 2.6 min) and low-freq (period 14 min) [2]. While studying other species of fungus, *Ganoderma resinaceum*, we found that most common width of an electrical potential spike is 5-8 min [4]. In both species of fungi we observed bursts of spiking in the trains of the spike similar to that observed in central nervous system [25, 42]. Whilst the similarly could be just phenomenological this indicates a possibility that mycelium networks transform information via interaction of spikes and trains of spikes in manner homologous to neurons. First evidence has been obtained that indeed fungi respond to mechanical, chemical and optical stimulation by changing pattern of its electrically activity and, in many cases, modifying characteristics of their spike trains [7, 8]. There is also evidence of electrical current participation in the interactions between mycelium and plant roots during formation of mycorrhiza [17]. In [30] we compared complexity measures of the fungal spiking train and sample text in European languages and found that the ’fungal language’ exceeds the European languages in morphological complexity.

**Figure 1:**
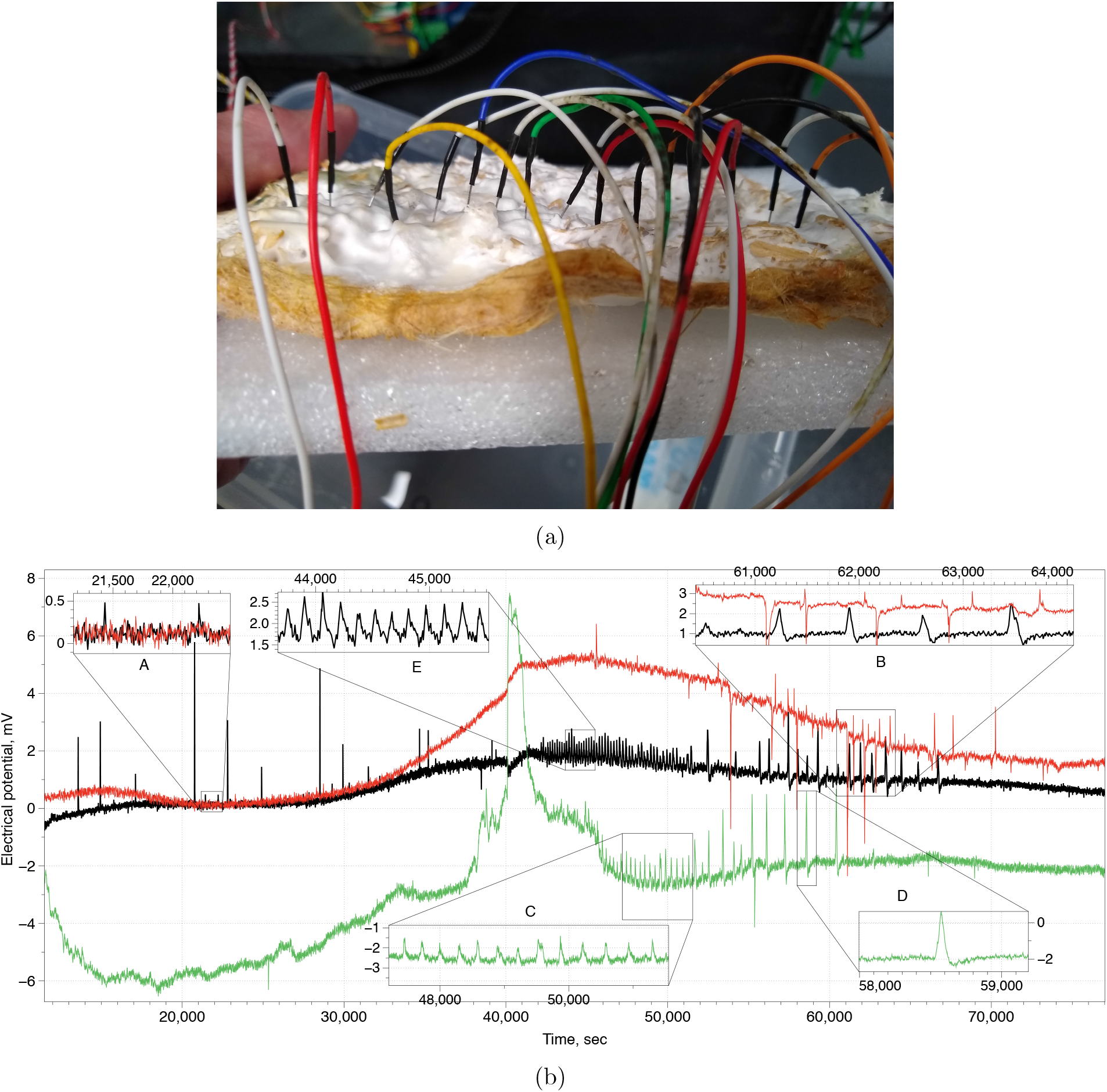
Recording electrical activity of fungi. (a) Setup with an array of differential electrodes pairs. (b) A variety of patterns of spike trains.

In [3] we recorded extracellular electrical activity of four species of fungi. We speculated that fungal electrical activity is a manifestation of the information communicated between distant parts of the fungal colonies and the information is encoded into trains of electrical potential spikes. We attempted to uncover key linguistic phenomena of the proposed fungal language. We found that distributions of lengths of spike trains, measured in a number of spikes, follow the distribution of word lengths in human languages. The size of fungal lexicon can be up to 50 words, however the core lexicon of most frequently used words does not exceed 15-20 words. Species *Schizophyllum commune* and *Omphalotus nidiformis* have largest lexicon while species *Cordyceps militaris* and *Flammulina velutipes* have less extensive one. Depending on the threshold of spikes grouping into words, average word length varies from 3.3 (*O. nidiformis*) to 8.9 (*C. militaris*). A fungal word length averaged over four species and two methods of spike grouping is 5.97 which is of the same range as an average word length in some human languages, e.g. 4.8 in English and 6 in Russian.

General anaesthetics in mammals causes reduction of neural fluctuation intensity, shift of electrical activity to a lower frequency spectrum, depression of firing rates, which are also reflected in a decrease in the spectral entropy of the electroencephalogram as the patient transits from the conscious to the unconscious state [56, 36, 38, 76]. In words, a rich spiking activity is a manifestation of consciousness, and reduced activity of unconsciousness.

In [5] we demonstrated that the electrical activity of the fungus *Pleurotus ostreatus* is a reliable indicator of the fungi anaesthesia. When exposed to a chloroform vapour the mycelium reduces frequency and amplitude of its spiking and, in most cases, cease to produce any electrical activity exceeding the noise level (Fig. 2). When the chloroform vapour is eliminated from the mycelium enclosure the mycelium electrical activity restores to a level similar to that before anaesthesia.

**Figure 2:**
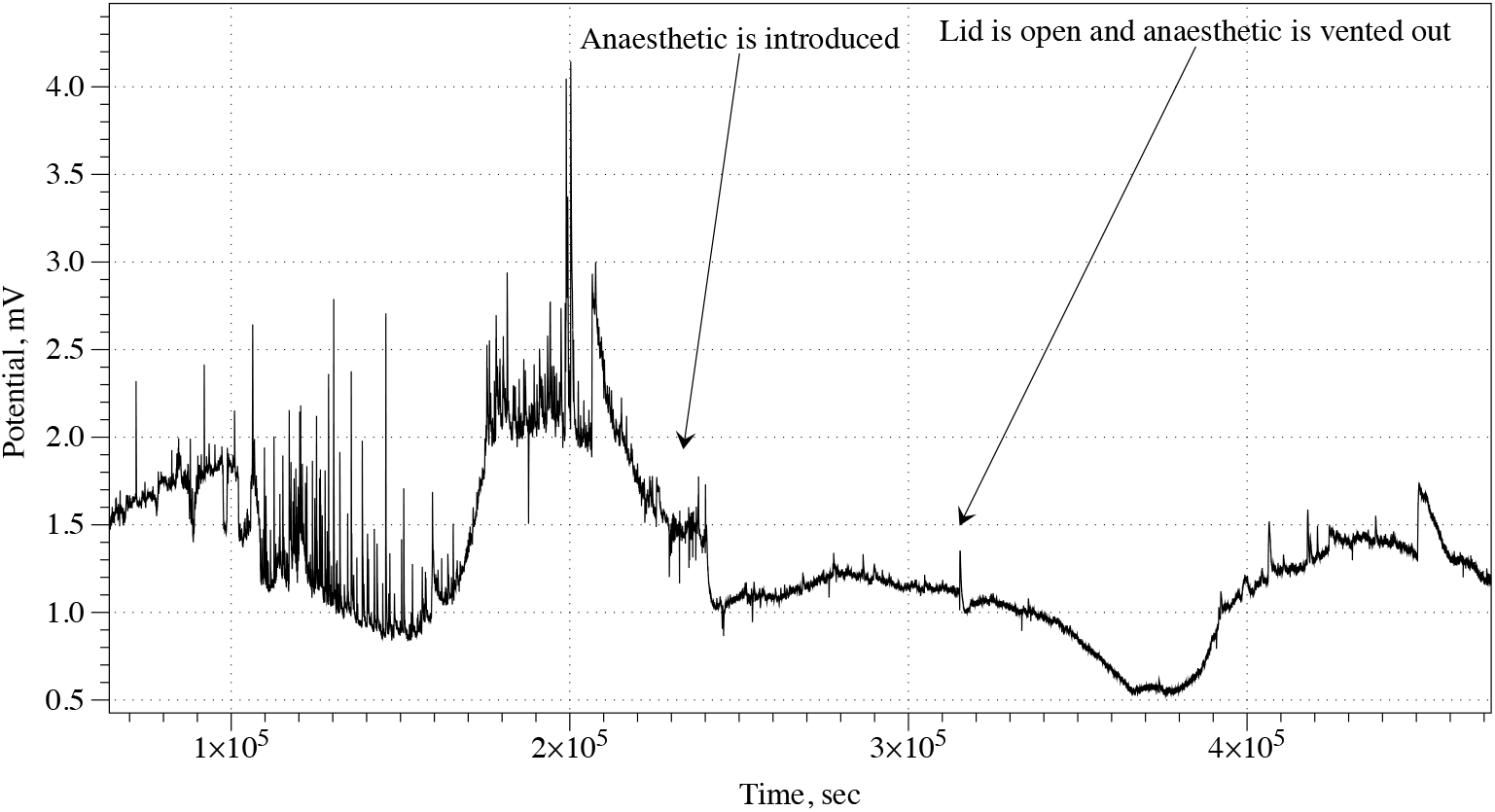
Reduction of spiking activity of *Pleurotus ostreatus* under influence of chloroform.

To summarise, in experimental laboratory studies of electrical activity of fungi we demonstrated that fungi produce neuron-like bursts of spikes which are affected by general anaesthetics. These phenomena indicate that fungi can posses the same degree of consciousness as creatures with central nervous system do.

## 3. Fungal cognition

Once accepting the unity of such a big fungal biological structure as a single living entity, we need to face a second challenge, that is, the anthropocentric bias [79] which sees consciousness as an exclusively human capacity. This latter is the main cause for the lack of interest in cognition and intelligence in minimal living systems. At this very moment we can affirm that intelligence is a property extended across all living taxa, and that we can even talk about minimal consciousness, starting at microbial level [54, 43, 44, 45, 51, 52, 16, 11]. The functional requests that make possible the existence of such huge fungal colony are beyond the simple or automated addition of neighbouring cells, but require a level of cooperation and informational communication that make necessary to ask for a mechanism that makes possible all these processes, could it be a form of consciousness? [53]. From a phylogenetic perspective fungi provide the mechanisms for the existence of plant synapses [11], a fundamental aspect for enabling plants complex information processing.

Our departure point is naturalistic and follows a simple idea: the biological explanations which can be identified using a functionalist approach supervene on chemical mechanisms; consequently, any approach to the emergence of informational minds must rely on such embodied factors. On the other hand, social interactions modify this process, forcing us to consider the emergence of mind as the coupling between single individual units and collective behaviour.

Our approach to the study of fungal minds is not a panpsychist one (attributing sentience to matter), but is based on an informational processing model in which we identify the fungal self-awareness mechanisms which provide an empirical foundation for the study of fungal minds. Two questions are orienting our study: are fungi sentient? and…if so, could we talk about fungal collective consciousness?

From mycorrhizal relations, we know that fungi interact with plants roots and allow the existence of mycorrhizal networks, used by plants to share or transport carbon, phosphorus, nitrogen, water, defense compounds, or allelochemicals. Thanks to this network, plants regulate better their survival, growth, and defence strategies. Such symbiotic relationship provides fungi carbohydrates, which are used metabolically to generate energy or to expand their hyphal networks, generating therefore the collective mycelium. And such mycelium can be considered the superstructure from which the collective fungal consciousness emerges. As magisterially described by Stamets [77] (page 4):

> The mycelium is an exposed sentient membrane, aware and responsive to changes in its environment. As hikers, deer, or insects walk across these sensitive filamentous nets, they leave impressions, and mycelia sense and respond to these movements. A complex and resourceful structure for sharing information, mycelium can adapt and evolve through the ever-changing forces of nature These sensitive mycelial membranes act as a collective fungal consciousness.

From an evolutionary point of view, mycelia are a clear example of cooperation, but also are used as a cheating mechanism in relation to host plants [22]. Cheating, besides, decreases when high genetic relatedness exists, a key point for sustaining multicellular cooperation in fungi [15]. We’ve considered possible cheating actions of fungi towards their mycorrhizal hosts, but how can they form collective living forms without cheating among themselves? The answer is related to the concept of allorecognition. The ability to distinguish self from non-self is beneficial not only for self-preservation purposes [69] but also for protecting the body from external menaces, like somatic parasitism [28] [66]. We’ve seen how fungi are able to distinguish between themselves and others, and how several mechanisms allow them to work in colonies, to establish symbiotic or parasitic relationships with other living systems. Their biochemistry allows them to adapt their actions to the informational variations of the surrounding conditions, and requires a cognitive system able to adaptively manage such actions.

## 4. Consciousness as self-cognition

When observing cognitive and intelligent behaviour (adequate decision making, learning, problem solving) of fungi we may ask whether some kind of consciousness enables their goal-directed behaviour, where consciousness is the ability to make sense of the present situation. One can search for the consciousness and its markers starting with humans and investigate its evolutionary origins in other living organisms. Comparing humans with simpler living organisms it might be useful to make the distinction between primary and higher order consciousness. With minimal modifications we can adapt the notion of “primary consciousness in humans to describe “primary consciousness” (that corresponds to “basal cognition” in other organisms including fungi and even unicellular organisms.

Over the centuries, consciousness has been a puzzling phenomenon despite all the efforts of the scientists who tried to unveil its mysteries. Plant cognition and intelligence has been a matter of study for hundreds of years, and still there is an ongoing controversy on its definition and functional extent [27]. There is a strong resistance and reluctance to acknowledge intelligence and cognitive capacities (including degree of consciousness) in other living beings.

Fungi can store information and recall it [32]. Fungal memories are procedural, what is associated with anoetic consciousness. Fungi show self-nonself recognition patterns. Fungi can navigate and solve mazes looking for a bait. Fungi perceive their environment guiding themselves to sources of light or higher oxygen concentrations. Self-nonself recognition, synchronous perception of light, nutrients, gravity, or gas, moisture or other chemical gradients, only involves response to internal or external stimuli, thus these cannot be quoted to support the affirmation that fungi are conscious or self-aware. Such behaviours can be also encoded in a computer, which will respond accordingly to the instructed parameters.

Fungus has a role in the managing of such aneuronal consciousness, as has been observed on tree colonies[34]. The fundamental part of our debate is to show that living systems without nervous systems (which we usually regard as necessary for self-awareness or intelligence) are able to perform tasks that we usually would ascribe to conscious systems, and that consciousness is ubiquitous[81].

## 5. Biosensing for Data Integrating: the Mechanistic Path to Fungi Consciousness

The first aim of consciousness, from a pragmatic point of view is that of collecting and combining information at some specific level of detail in order to take a decision for action. Up to now the main focus on consciousness studies has been focused on high level cognitive processes that follow a top-down hierarchical structure. Using this model, fungi should be automatically discarded as suitable living systems which could show consciousness. Instead of it our approach to the notion of consciousness will follow a bottom up approach, from basic data to its ulterior processing and the possible conscious decisions. We will present this case from a context situation: Mycelium typically is just under the *humus* (the soil cover, i.e. a mix of leaves, needles from pines, fallen branches etc). Thus when we walk in the forest mycelium “knows” by mechanical stimulation and stretch-activated receptors [21] that we are walking there. This mechanoception process is shared by several living systems, including plants [58].

Consider for example calcium signal transduction [49]: it has been proven that in fungi this signalling pathway has an essential role in the survival of fungi [88], as well as mediate stress responses, or promote virulence [61]. Mechanosensitive channels can also be important for mating, as we see in *Neurosopora Crassa* [46]. The existence of such mechanosensitive receptors in fungi make possible to extend some cognitive properties we have already clearly defined in plants or mammals to fungi. Consider for example the purinergic signalling [1]. Furthermore, sensing capabilities so far described include also nutrient sensing (glucose, nitrogen) and general chemophysical sensing (pH, temperature, light, gravity, electric field) [21]. Are those mechanisms a sufficient basis for the grounding of the following questions: What does mycelium feel about this? Can the mycelium trace our movement? Can the mycelium predict that we are approaching fruiting bodies (mushrooms)?

### 5.1. Integration data in fungi systems

All cognitive systems display mechanisms for using captured information (an active process, not just a passive one) and to decide output actions. Due to the multimodal nature of data, this process implies an integration and meaning hierarchies. The properties of fungi in data integration are shown in Tab. 1.

**Table 1:** Different cognitive tasks performed by fungi.

| Fungal abilities | Evidence |
| --- | --- |
| Decision-making and spatial recognition | Fungi use an elaborate growth and space-searching strategy comprising two algorithmic subsets: long-range directional memory of individual hyphae and inducement of branching by physical obstruction [33]. The human pathogenic fungus *Candida albicans* is able to reorient thigmotropically its hyphae to find entry points into hosts [78]. |
| Short-term memory and learning | Fungi exposed to a milder temperature (priming) stress perform better when exposed to a potentially damaging second heat (triggering) stress. The priming state in filamentous fungi dissipates over time: memory of the initial priming stress event for a period of time of at least 24 h [10]. Mycelium of *Phanerochaete velutina* remembered the location in which a bait was placed in a previous test, growing towards the same direction in a new empty tray [32]. Saturating light stimulus habituates *Phycomyces sporangiophores* to a light stimulus but not to an avoidance stimulus [64]. |
| Long-distance communication | Vacuole-mediated long-distance transporting systems support mycelial foraging and long-distance communication in saprophytes and mycorrhizal fungi [82, 87]. |
| Photo-tropism | Fungi show a rich spectrum of responses to light. They sense near-ultraviolet, blue, green, red and far-red light using up to 11 photoreceptors [24, 48, 65, 89]. |
| Gravi-tropism | Gravitropism, as well as thigmotropism, is the strongest tropisms of fungi. This tropism is well studied and documented [59, 26] |
| Chemo-tropism and chemical sensitivity | Fungi are able to detect sources of nutrients and grow towards them (foraging), in a similar fashion, a fungus would react against a harmful chemical trace (e.g. toxic metals) by growing towards the opposite direction [18, 31]. Fungal colonies communicate through volatile compounds [40, 12, 39]. |
| Sensing touch and weight | Thigmo-based responses, include thigmo differentiation, thigmonasty, thigmotropism [67, 19, 9, 8, 6]. |
| Self vs. non-self recognition | Fungi possesses incompatibility loci and a genetic difference at any one of them is sufficient to trigger destruction of the mixed cell: in most fungal species, pairs of isolates taken at random are generally incompatible [66]. |
| Fighting behaviour | Several species of fungi are capable for capturing and consuming nematodes [13, 50]. Antagonistic interactions between wood decay basidiomycetes show combative hierarchies with different species possessing different combinations of attack and defence traits [37]. |
| Trade behaviour | The nutrient exchange mutualism between arbuscular mycorrhizal fungi (AMFs) and their host plants qualifies as a biological market [62, 41, 74, 73]. |
| Manipulating other organisms | Fungi evolved elaborate tactics, techniques and molecular mechanisms to control other organisms, from attracting and paralysing nematodes to programming insect behaviour and death [86, 72, 55] |

### 5.2. The Self: from the genetic view to the perceptual and processing ones

It is necessary to say that the search for published scientific papers on the topics about “fungal cognition” offers us zero results. The only connections between fungi and cognition are related most of times to the cognitive impact for humans who are in contact with fungi. But, of course, fungi do perform cognitive tasks, being self perception one of the most important.The most basic notion of self identification is related to the delimitation of the structural elements that belong or not to one system. In this sense, fungi have shown that their hyphae are able to discriminate self/non-self and that use this skill to decide to fuse themselves with other genetically compatible hyphae, thanks to anastomosis [70]. The mycelial networks of mycorrhizal fungi can also recognise correctly the roots of their hosts from those of other surrounding non-hosts. Even different mycorrhizas can coexist but never fuse together. There is a second mechanism implied in self recognition that is omitted in most of current studies: alarmones [83]. These are regulatory molecules used to communicate exclusively among single cells (not as host). Following [83], pages 13-14, we notice that a static organism, like a filamentous fungi can live very long and move by hyphal growth, although it is physically constrained and must withstand the onslaught of all potential genetic parasites they will encounter in their long life. The skill of self-identification and colonial identity is therefore fundamental for several purposes (such as mating control). And some fungi even use retro-parasites for their own development.

## 6. Discussion

We presented experimental laboratory and philosophical studies of the fungal states of mind. We considered several aspects of fungal cognition and provided arguments supporting existence of fungal consciousness. We raised many more questions than we provided answers. The new field of fungal consciousness is opening in front of us. Let us discuss directions of future studies. How could we make sure (in wording) that the fungal behaviour is not mechanical (automatic) responses but that it holds intention? Abstraction, creativity, judgement, are characteristics of human consciousness. Are fungi able of performing these? It might be useful to make the distinction between “primary” and “higher order” consciousness.

Do fungi have holistic states of mind? Do they combine/modify such states? How many fungal states of mind could be described? Do fungi create specific relational contexts? Or are they, on the contrary, not capable of having holistic states of mind, ajust following completely prefixed patterns? At which extent can we include fungi affects into such cognitive processing? Fungal chemotaxis could be part of such proto-emotional states.

A deeper consciousness state allows us to understand and accept sacrificing our “self“ for higher purposes (martyrs, heroes). Nevertheless we must recognise that human individuals (and animals as well) are genetically different from each other, with the only exception of twins, while the same could not apply to fungi. This in part supports the observation that fungi sacrifice their bodies for the sake of their propagation. Another thought is about mortality: though it is, particularly in the western civilisation, uncommon for people to live everyday life with a constant thought of being mortal, and pursue choices that quite often go in the opposite direction, as if the individuals are immortal, it is unquestionably difficult for humans to figure out how the world would appear for immortal beings, like certain fungi. An immortal, or even extremely old consciousness, would be able to develop perhaps an intelligence out of reach for us, pursuing objectives that might seem unreasonable, for our limited perception. Perception of space and time, causality, are all aspects that we consider our unquestionable bottom line. But given the peculiarity of fungi morphology and degree of connection, we may imagine how radically different computational schemes are embedded into a fungal consciousness. For example, rather than 3-dimensional visual perception, holographic perception might be possible, considering the quasi-flat distribution of mycelia and its mechanoceptive reconstruction of moving objects (animals) at the upper boundary layer. Non-causal consciousness might arise from this specific perception framework, eventually hindering the time perception. All of these remain open questions for further investigations.

## 7. Acknowledgement

AA and NP were supported by the funding from the European Union’s Horizon 2020 research and innovation programme FET OPEN “Challenging current thinking” under grant agreement No 858132.

